# The eXplainable Artificial Intelligence (XAI) Triad: Models, Importances, and Significance at Scale

**DOI:** 10.64898/2025.12.06.692723

**Authors:** Ghislain Fievet, Julien Broséus, David Meyre, Sébastien Hergalant

**Affiliations:** NGERE - Nutrition-Génétique et Exposition aux Risques Environnementaux; CHRU Nancy - Centre Hospitalier Régional Universitaire de Nancy

## Abstract

In this study, we present a comprehensive evaluation framework for comparing various combinations of artificial intelligence (AI) methods in the context of explainable AI (XAI) for variable selection in experimental biological and biomedical data. Our goal was to assess the efficiency, computational cost, and accuracy of different method combinations across six simulated scenarios, each replicated ten times. These scenarios encompass various classification and regression complexities, including variance differences, bimodal distributions, eXclusive-OR (XOR) interactions, concentric circles, and nonlinear relationships such as parabolic and sinusoidal functions.

We tested several machine learning algorithms, including Decision Trees (DT), Random Forests (RF), Support Vector Machines (SVM), and Multi-Layer Perceptrons (MLP). We combined these models with diverse feature-importance methods such as Gini importance, accuracy decrease, SHAP values (Shapley Additive exPlanations), and Olden’s method. We further applied significance-thresholding approaches, namely PIMP (Permutation IMPortance), mProbes, and the novel simThresh developed for this study. Additionally, we explored different dataset sizes to evaluate the scalability of these methods. Our analysis revealed substantial differences in computational demands, ranging from very rapid evaluations (e.g., DT combined with Gini importance and simThresh averaging 0.15 seconds) to extensive computations (e.g., MLP combined with SHAP and PIMP exceeding 7 hours).

Among the tested combinations, RF/Accuracy/PIMP achieved the best overall performance, successfully identifying 59 out of 60 replicates in our benchmark study. However, this approach raises concerns regarding its scalability when applied to large-scale omics datasets in real-world settings due to its computational demands. In contrast, Decision Tree or Random Forest models using Gini and simThresh criteria ranked second, with 50 out of 60 detections. While less accurate, these methods require fewer computational resources, making them more promising candidates for scalable applications in omics data analysis. The proposed evaluation framework thus serves as a valuable tool for method selection, particularly relevant when dealing with large-scale omics datasets where computational resources and accuracy are both critical considerations.

## Introduction

In recent years, researchers have increasingly turned to XAI techniques to analyze biomedical datasets. A 2023 systematic mapping study identified over 400 papers applying XAI in genomics, transcriptomics, proteomics, and metabolomics, with feature importance (FI), SHAP [1] being among the most commonly used [2]. This surge reflects the growing demand for transparency in machine learning (ML) models for biomedicine, as XAI promises to identify which molecular features or patterns drive predictive outcomes.

However, different combinations of ML models and XAI methods can vary greatly in their ability to uncover specific types of relationships in data. A classic example is a XOR-type interaction, where two features jointly determine an outcome but each feature alone appears uninformative. As an illustration, a recent evaluation of multiple XAI techniques on XOR-based data found that none perfectly recovered the true importance of both interacting features under all conditions [3].

In real-world omics research, new tools were recently developed to harness the discovery potential of XAI methods, which help interpret model predictions and uncover putative biomarkers. For example, the recent OmicsFootPrint platform integrated deep neural networks with SHAP to interpret multi-omics models of cancer, successfully pinpointing critical genes and proteins that drive disease subtype classifications [4]. Such applications demonstrate the promise of combining advanced ML with XAI: by attributing importance to genomic or proteomic features, researchers can generate hypotheses about key pathways or molecular drivers underlying a phenotype.

At the same time, there is growing awareness that biomedical researchers must use these XAI outputs cautiously, with an understanding of each method’s sensitivities and blind spots. Studies have noted that SHAP’s feature rankings can vary significantly with model architecture and random initialization, raising concerns about the consistency and trustworthiness of the explanations in deep learning frameworks. In one multi-omics study, the authors reported some difficulties in validating certain biological associations flagged by SHAP, acknowledging that inherent limitations of the Shapley-value approach might be responsible [5]. These challenges highlight the need for dedicated tools and evaluation strategies to assess not only the robustness and biological relevance of XAI-derived insights, but also the kinds of patterns such methods are truly capable of detecting.

A critical aspect of FI attribution is often overlooked: the statistical significance of the reported importances. Most studies rely on raw scores (e.g., SHAP values or Gini indices) without assessing whether these reflect true associations or result from random fluctuations in the data. However, several methods have been developed to address this issue, notably PIMP (Permutation Importance) [6] and mProbes [7], which estimate null distributions of importance values to derive p-values. However, these approaches remain underused and have not been systematically compared, especially in combination with modern XAI tools across diverse data structures.

To address this gap, we designed a benchmarking framework that evaluates the performance of multiple end-to-end pipelines for variable selection. These pipelines combine:

- several machine learning models, such as DT, RF, SVM and MLP.
- various feature attribution methods, SHAP, Gini importance, accuracy and Olden score.
- diverse statistical thresholding techniques to select significant features, PIMP, mProbes and simThresh, a novel method we introduce here based on the addition of a specifically generated feature used as a reference threshold.

We applied these combinations to a set of six synthetic scenarios that reflect different classification and regression situations. By doing so, we aim to identify robust model + importance + significance configurations that offer reliable signal detection while maintaining computational efficiency, a first step towards larger dataset analyses.

## Material and methods

### Synthetic data scenarios

We designed four classification and two regression scenarios that represent signal patterns typically undetectable by standard univariate methods. Specifically, the classification scenarios were constructed to be invisible to traditional t-tests or Wilcoxon tests, while the regression scenarios were not detectable via Pearson correlation or simple linear models.

Each classification dataset included 100 samples (50 per class), with a background of 10 Gaussian noise features, two features with a classical mean shift between classes (as internal positive controls), and one or two features specific to the scenario under evaluation. Each regression dataset consisted of 100 samples, 10 Gaussian noise features, 10 uniform noise features, and one feature encoding the signal to detect.

- **Scenario 1, Variance Difference** (Classification) : Two classes are drawn from the same mean but with distinct variances on one feature.
- **Scenario 2, Bimodal Distribution** (Classification) : Class separation is created using a feature with different bimodal distributions. This scenario emulates the case of hidden subgroups or mixed populations (e.g., disease subtypes) and introduces challenges when the means are not good discriminants.
- **Scenario 3, XOR Interaction** (Classification) : The outcome depends on an XOR logical interaction between two features, each feature is individually non-informative but jointly predictive. This classic nonlinear interaction tests whether methods can detect feature dependencies beyond marginal effects.
- **Scenario 4, Concentric Circles** (Classification) : Class labels are determined by how close a point is to the center, forming two circular groups. This scenario tests whether methods can detect round-shaped patterns rather than straight-line separations.
- **Scenario 5, Parabolic Relationship** (Regression) : The target variable follows a parabolic curve, where values increase and then decrease, or vice versa. This tests whether methods can detect simple nonlinear patterns instead of just straight-line relationships.
- **Scenario 6, Rising Sinusoid** (Regression) : The response variable follows a sinusoidal pattern with increasing amplitude as a function of one input feature. This case reflects complex oscillatory patterns.

Each scenario was independently generated ten times to assess robustness, using different random draws in each replicate. All synthetic datasets were generated using the *genScenario* function from the *XAItest* package. This function reproduces the six benchmark scenarios and includes options to modify the number of samples and noise features, making it suitable for scalability testing of the methods.

### Machine Learning Models

We evaluated four commonly used supervised machine learning algorithms, selected to represent diverse model families and learning strategies:

- **Decision Tree** : A simple and interpretable tree-based model that recursively partitions the feature space to minimize impurity. We used the default *rpart* implementation in *R*.
- **Random Forest** : An ensemble of decision trees trained on bootstrapped samples and random feature subsets. We used the *randomForest* package with default parameters, setting the number of trees to 500.
- **Support Vector Machine** : A classifier that finds the optimal separating hyperplane in high-dimensional space. We used both the linear (*vanilla*) and radial basis function (RBF) kernel variants from the *e1071* implementation.
- **Multi-Layer Perceptron** : A feedforward neural network composed of two hidden layers (64 and 32 neurons) with *ReLU* activations, implemented using the *keras R* interface. The network was trained using the Adam optimizer with 5000 epochs.

For DT, RF, and SVMs, we used default parameters without hyperparameter tuning, relying on their established robustness and simplicity.

For the MLP, which is more sensitive to architectural choices and prone to overfitting, we performed an initial 50/50 train/test split to select an architecture that yield stable performance without overfitting. This step ensured that the chosen network could learn relevant patterns while maintaining generalizability.

Once validated, these same model configurations were retrained on the full dataset to perform feature attribution. This approach is justified by the fact that our objective is not to evaluate generalization accuracy, but to test the ability of methods to identify relevant features. Using the full dataset avoids reducing statistical power and does not bias the results in our context. Unlike typical settings where test data must remain unseen to prevent optimistic bias, here the risk is inverted: overfitting would obscure rather than artificially enhance signal detection. If an explainer still successfully recovers the correct variables after training on all data, it strongly suggests that the detection is robust, not an artifact of overfitting.

### Feature Importance

After training each model on the ten replicates of the six scenario datasets, we computed a feature importance score for every input variable in each model. We applied four feature importance algorithms, selected based on their compatibility with the corresponding machine learning models :

- **SHAP** : SHAP values provide a game-theoretic approach to explain the output of any model by assigning a contribution score to each feature. We used SHAP with all models using the R package *kernelshap*.
- **Mean Accuracy Decrease** : This method estimates the importance of a feature by quantifying the reduction in predictive accuracy when its values are randomly permuted. It was applied to Random Forests, for which this metric is natively computed during training by the *randomForest* packages.
- **Gini Importance** : Based on the decrease in node impurity during training, Gini importance is a native feature ranking method available for tree-based models. The Gini index measures the probability of misclassifying a randomly chosen sample if it were labeled according to the class distribution in a node. A Gini value of 0 indicates perfect purity (all samples in the node belong to the same class), whereas a Gini value of 1 indicates maximum impurity (samples are uniformly distributed across classes). Gini importance sums the impurity decreases contributed by each feature across all splits. It was applied to Decision Trees and Random Forests, for which this metric is natively computed during training by the *rpart* and *randomForest* packages, respectively.
- **Olden’s Method** : Specific to neural networks, Olden’s method estimates feature importance by analyzing the contribution of input neurons through hidden layers to the output layer, based on the product of connection weights. This method was used exclusively with MLPs and was implemented using the *NeuralNetTools* package in R.

### Conversion into Statistical Significance

While raw feature importance scores provide a ranking of variables, they do not indicate whether a given importance value is statistically meaningful or could arise by chance. To address this, we applied three significance thresholding methods to convert feature importance values into p-values, allowing for a principled selection of relevant variables.

#### PIMP

The PIMP approach estimates the null distribution of feature importance by repeatedly permuting the target variable, retraining the machine learning model and recomputing importance scores. A p-value is then derived by comparing the observed importance to this null distribution. In our workflow, PIMP was applied with 100 permutations per replicate.

#### mProbes

The mProbes method adds a set of artificial “probe” features, generated from random noise, to the dataset. Because these probes have no relationship with the target, their importance values provide an empirical reference for the null distribution. Real variables are considered significant if their importance is greater than that of the probes. In our workflow, mProbes was applied with 10 probes and 100 permutations per replicate.

#### simThresh (proposed method)

We introduce a method called simThresh. Unlike PIMP and mProbes, simThresh does not convert feature importances into p-values, but instead defines a feature importance threshold above which the value is considered significant.

For **classification**, the approach is based on creating a simulated variable whose values follow class-specific distributions, with an adjustable difference between the classes. The goal is to find the difference δ between two distributions 𝒟 (*μ, σ*^2^) such that :

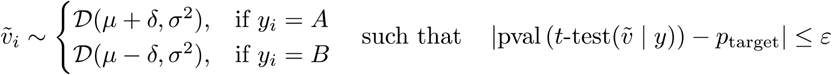

Where :

- 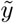 is the simulated variable
- *y* is the categorical variable
- 𝒟 (*μ, σ*^2^) is a distribution with mean *μ* and variance *σ*^2^
- *p* _*target*_ is the target p-value
- *ε* is the allowed tolerance on the p-value

A dichotomous search process is used: if the p-value is too high, the difference is increased; if it is too low, it is reduced.

For **regression**, the simulated variable is created by adding noise to the target variable, with a standard deviation adjusted to control the correlation p-value between the simulated variable and the target. The goal is to find the noise level *σ* from 𝒟 (*μ, σ*^2^) such that:

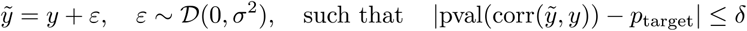

where :

- 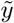 is the simulated variable
- *y* is the target variable
- 𝒟 (*μ, σ*^2^) is a distribution with mean *μ* and variance *σ*^2^
- *p* _*target*_ is the target p-value
- *δ* is the allowed p-value tolerance

This iterative process continues until the p-value reaches the predefined target within the specified tolerance. The feature importance associated with this calibrated simulated variable is then used as the significance threshold, against which the scores of real variables are compared. This approach provides scenario-specific significance control while keeping computational cost low: the dichotomous search phase requires only rapid t-tests or Wilcoxon tests, and the second phase requires training the ML model only once with the simulated variable included in the dataset, unlike PIMP or mProbes, which require multiple retrainings.

## Results

### Evaluation protocol

#### Removing trivial variables (two-stage evaluation)

To prevent models from “coasting” on trivial signals, we evaluated each pipeline twice: (i) on the full dataset and (ii) on a reduced dataset obtained by removing variables that classical univariate tests (t-test or Wilcoxon test) can already detect via a simple mean shift. This procedure isolates the method’s ability to recover the harder, scenario-specific signals rather than relying on trivial features. Comparing results on full *versus* reduced sets clarifies whether a pipeline truly targets the intended patterns.

#### Success criteria

We defined four hierarchical success criteria spanning full/reduced datasets and threshold/ranking strategies. A method “succeeds” if it exactly retrieves the ground-truth markers under one of the following :

1. 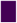 markers are exactly those with p-value < 0.05 on the full dataset
2. 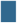 markers are exactly those with p-value < 0.05 on the reduced dataset
3. 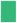 markers are exactly those at the top of the feature-importance ranking on the full dataset
4. 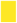 markers are exactly those at the top of the feature-importance ranking on the reduced dataset

These four criteria are ranked from the most stringent to the most permissive: meeting criterion (1.) means that the method can detect the correct variables with statistical significance, even in the presence of trivial variables. Criterion (2.) relaxes this requirement by removing these trivial variables, identifiable by a t-test, which may help some methods focus on the more complex variables. Criteria (3.) and (4.), based on ranking rather than p-values, are more flexible but still provide valuable insight into a method’s ability to correctly prioritize the relevant variables correctly. Together, these four criteria enable a progressive and nuanced assessment of the performance of the tested methods.

### Computational cost

#### Why time matters

An important component of our evaluation is the computation time associated with each tested combination of model, feature importance method, and statistical significance approach. Computation time is often a problem when applying XAI to large models and datasets. Several studies note that while SHAP provides detailed explanations, it is computationally expensive and can become impractical for large-scale analyses, which limits its scalability [8], [9], [10], [11], [12].

Depending on the methods used, total runtime can vary from a single, rapid training followed by one inference, to computationally intensive workflows involving up to 100 repeated trainings, potentially slow, and 100 inferences per training. For example, the combination Decision Tree + Gini importance + simThresh is computationally efficient : the decision tree model trains rapidly, Gini importance values are available immediately after training, and simThresh determines statistical significance without requiring any model retraining or additional inferences. In contrast, the combination MLP + SHAP + PIMP (or mProbes) is considerably more time consuming. The MLP requires longer training, in our experiments, we performed 100 inferences to estimate FI with SHAP. When coupled with PIMP or mProbes, this process must be repeated 100 times to build a null distribution, resulting in a combinatorial increase in processing time.

While these runtimes remain feasible for our simulated datasets of 100 samples and between 13 and 21 variables, the cost escalates rapidly for larger, real-world datasets. For instance, scRNA-seq experiments often measure 30,000 genes across tens of thousands of cells. DNA methylation arrays can reach 850,000 variables [13], and proteomics or bulk RNA-seq datasets typically range from 2,000 to 20,000 variables. At such scales, approaches requiring repeated retraining or multiple inference passes become computationally prohibitive.

Given the substantial variation in complexity across method combinations, and the large size differences between simulated and real datasets, systematic assessment of computation time is critical when selecting practical and scalable analysis pipelines.

## Results

### Computational cost and scalability analysis

Grouping combinations by whether they used MLP, SHAP, and/or simThresh yielded stark contrasts. The fastest group was simThresh + no-MLP + no-SHAP with a mean of 0.15 s, while the slowest was PIMP/mProbes + SHAP + MLP with 25,953 s (~7 h), a 173,000× ratio. Observed rules-of-thumb :

- DT/RF/SVM < MLP (model training cost)
- Gini / accuracy / Olden < SHAP (importance cost)
- simThresh < PIMP, mProbes (significance cost)

These contrasts reflect the long training time of MLPs and the heavy computations required by SHAP, PIMP, and mProbes.

To further explore scalability, we evaluated execution time as a function of dataset size across all model/importance/significance combinations (Figure 3). While the previous experiments focused on fixed datasets of 100 samples and between 13 and 21 features, here we varied the number of samples from 10 to 10,000, with the number of features set to twice the sample size. This design emphasizes how computation costs grow with increasing data dimensionality. The results show clear differences between methods: tree-based models with Gini importance or accuracy decrease and simThresh remained fast even for large datasets, while combinations involving MLPs, and resampling-based algorithms such as SHAP, PIMP and mProbes quickly became prohibitively time consuming, which shows that these approaches are not scalable.

**Figure 1.**
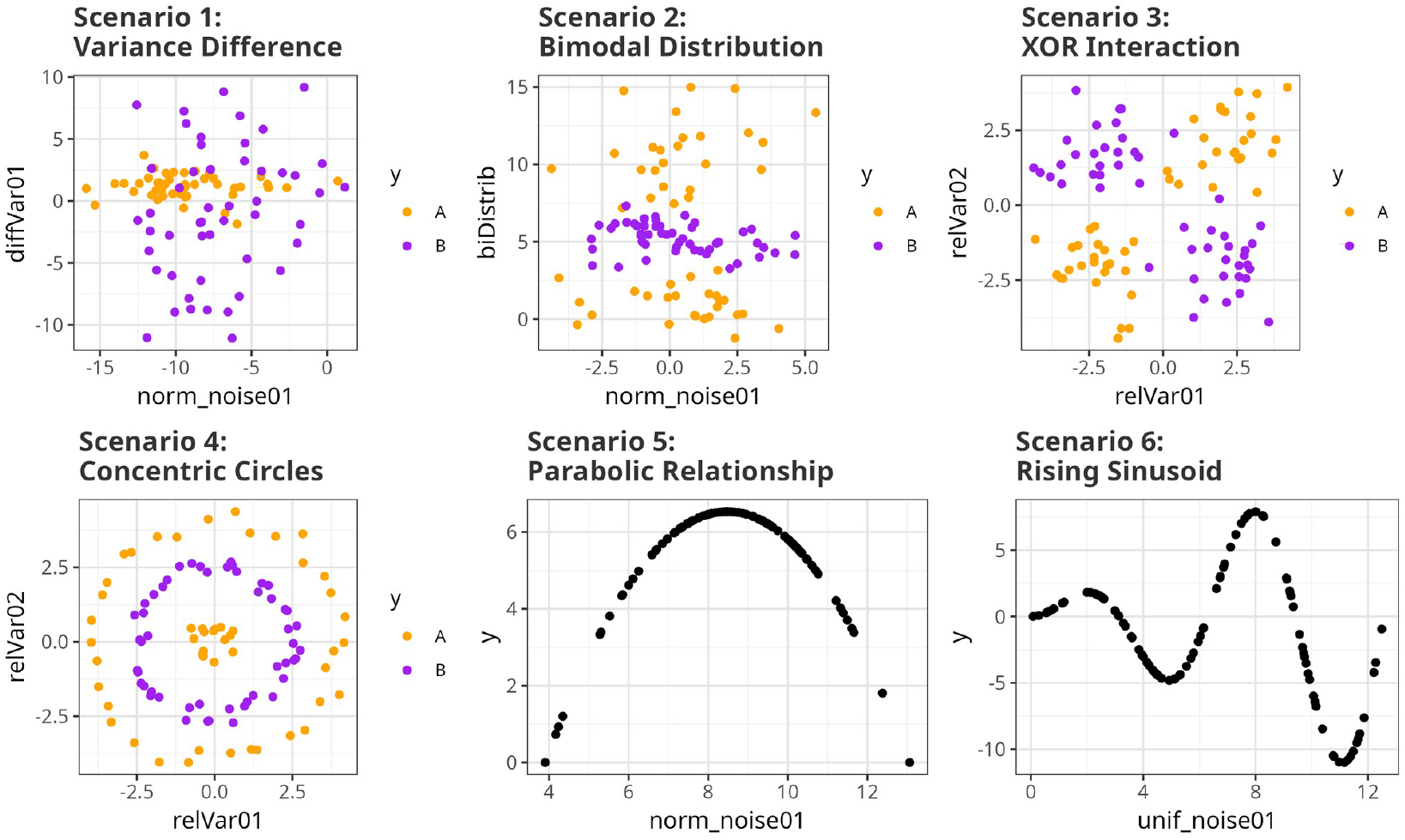
Overview of the six simulated scenarios used to benchmark XAI pipelines. The four classification tasks require separating classes based on: (1) different variances with identical means; (2) a bimodal versus a unimodal distribution; (3) a logical XOR interaction between two features; and (4) a concentric circle pattern. The two regression tasks involve fitting (5) a parabolic relationship and (6) a rising sinusoidal curve. These scenarios were specifically designed to be challenging for traditional statistical tests but solvable by machine learning models, thereby assessing the discovery power of different XAI methods.

**Figure 2.**
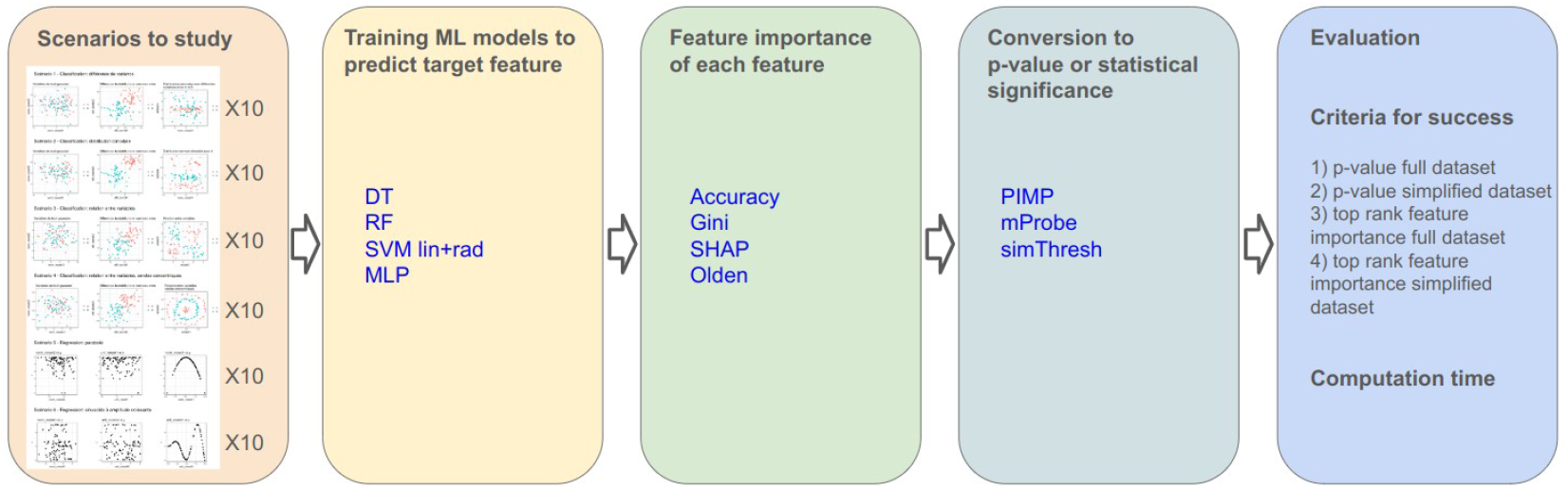
The triad model + importance + significance workflow.

**Figure 3.**
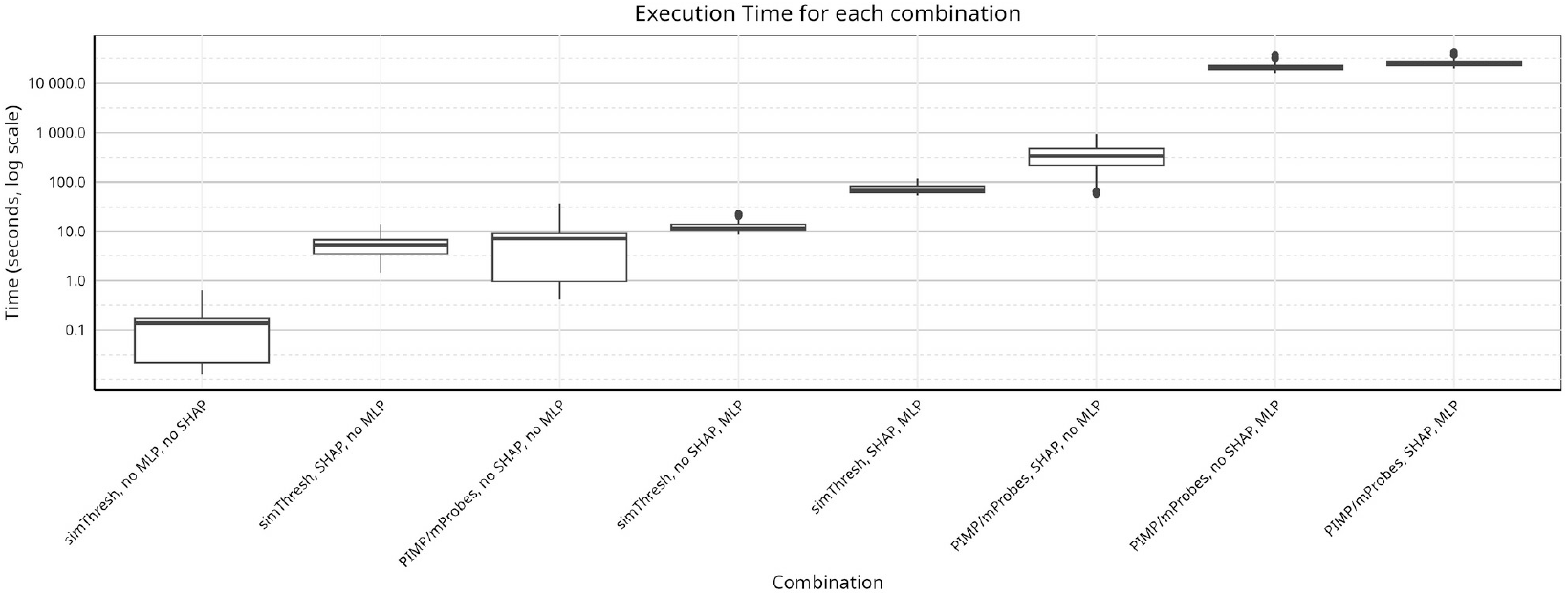
Comparison of execution times across different categories of XAI pipelines. The box plots summarize the total computation time (y-axis, log scale) for method combinations, computed over 10 replicates across six scenarios, and grouped by their core components (with or without MLP, with or without SHAP, and with or without simThresh). The results reveal a significant disparity in computational cost between the different approaches.

**Figure 4.**
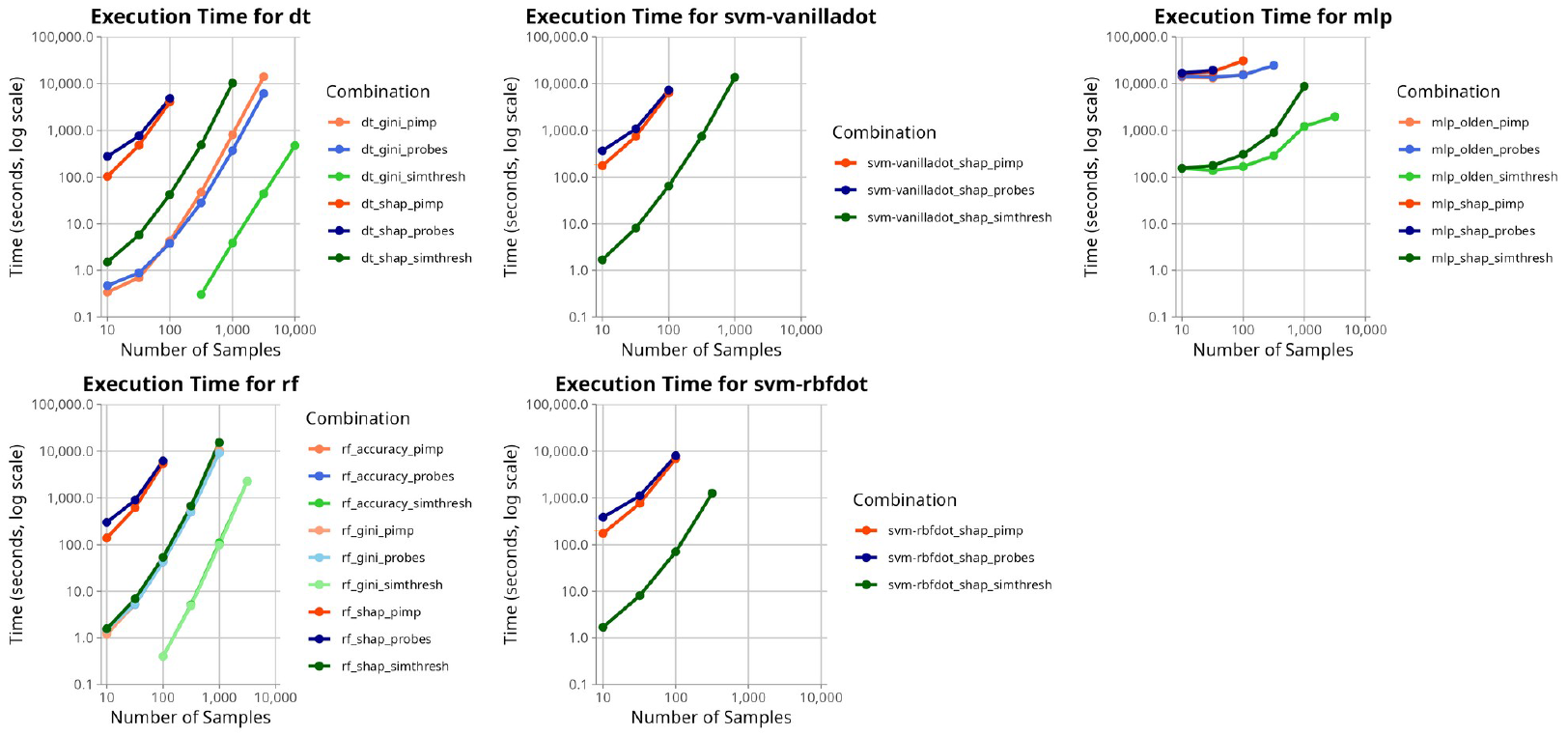
Execution time vs. dataset size for each model. For each x-axis value n, the dataset has n samples and 2*n features (matrix n×2n). Y-axis: seconds (log scale).

**Figure 5.**
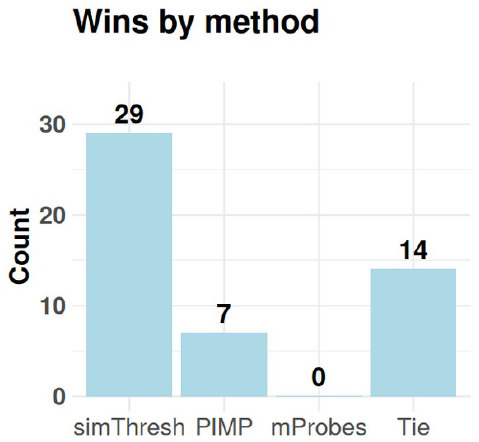
Comparison of significance methods based on the number of configurations where they achieved the highest detection rate of relevant features across scenarios, machine learning models, and feature importance methods.

### Detection Capabilities Across Models and Scenarios

Because simThresh yields a threshold exceedance rather than per-feature p-values, rank-based success criteria (our 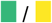 categories) do not apply to simThresh tables. In regression scenarios (5–6), all methods lack 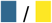 entries because each dataset contains only one variable of interest, and thus there is no need to consider any reduced dataset.

### SHAP rarely beats native/alternative FI

A useful way to evaluate the relative performance of feature importance methods is to organize them into pairwise “battles”, comparing SHAP against the most relevant alternative for each model type (Supp. Table 1). For decision trees, SHAP can be directly compared to Gini importance; for random forests, to both Gini and mean accuracy decrease; and for multilayer perceptrons, to Olden’s score. For SVM, SHAP is the only available FI method in our study.

When aggregating results across all non-tie conditions, SHAP outperformed its comparator only 4 times out of 31 . Even in scenarios where SHAP did identify the correct features, its computational cost was significantly higher than that of the competing methods. Together, these findings suggest that while SHAP remains a valuable, model-agnostic tool, it rarely offers an advantage in detection power over simpler, model-native alternatives, which are often both faster and more effective, especially in large-scale omics contexts.

### simThresh vs. PIMP vs. mProbes

To systematically compare the three significance strategies, we examined, for each scenario, machine learning algorithm, and feature importance method, which significance approach detected the relevant features most effectively (Supp. Table 2). The results show that simThresh consistently outperformed both PIMP and mProbes, being the best method in 29 out of 36 non-tie configurations, compared to 7 for PIMP, none for mProbes, and 14 ties overall.

### Detection performance varies across scenarios

Scenario 2, corresponding to the bimodal distribution, is, for example, detected with an average of 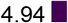 detections out of 10 (considering only simThresh and PIMP), and 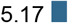 out of 10 replicates per combination. Scenario 3, corresponding to the XOR interaction, is detected much less frequently, with an average of 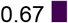 out of 10 and 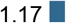 out of 10. This allows us to establish a ranking of scenarios according to their detection difficulty.

Among our scenarios, the most easily detected are the regression ones, i.e., Scenarios 5 and 6, with detection rates of 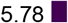 and 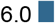 out of 10, respectively. The most difficult scenarios to detect are those involving interactions between multiple variables, XOR (Scenario 3) and concentric circles (Scenario 4), with average scores of 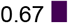 and 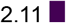 out of 10, respectively. Scenarios based on variance difference (Scenario 1) and bimodal distribution (Scenario 2) show intermediate difficulty, with average scores of 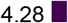 and 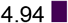 out of 10, respectively.

**Table 1:**
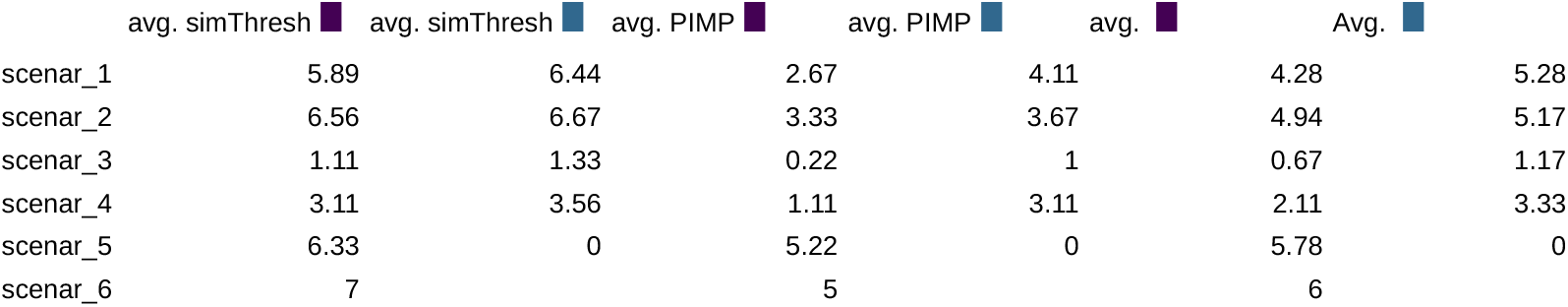
Comparison of detection rates for simThresh and PIMP across six benchmark scenarios.

|  | avg. *simThresh* | avg. *simThresh* | avg. *PIMP* | avg. *PIMP* | avg. | Avg. |
| --- | --- | --- | --- | --- | --- | --- |
| scenar_1 | 5.89 | 6.44 | 2.67 | 4.11 | 4.28 | 5.28 |
| scenar_2 | 6.56 | 6.67 | 3.33 | 3.67 | 4.94 | 5.17 |
| scenar_3 | 1.11 | 1.33 | 0.22 | 1 | 0.67 | 1.17 |
| scenar_4 | 3.11 | 3.56 | 1.11 | 3.11 | 2.11 | 3.33 |
| scenar_5 | 6.33 | 0 | 5.22 | 0 | 5.78 | 0 |
| scenar_6 | 7 |  | 5 |  | 6 |  |

### Balancing detection power and computational cost

The results show how each scenario favors different combinations of model, feature-importance method, and significance thresholding, with some configurations succeeding under specific data structures while others fail (Fig. 6A). When these outcomes are aggregated across all scenarios, a clear ranking emerges, revealing which approaches provide the most consistent detection of the relevant variables (Fig. 6B).

**Figure 6.**
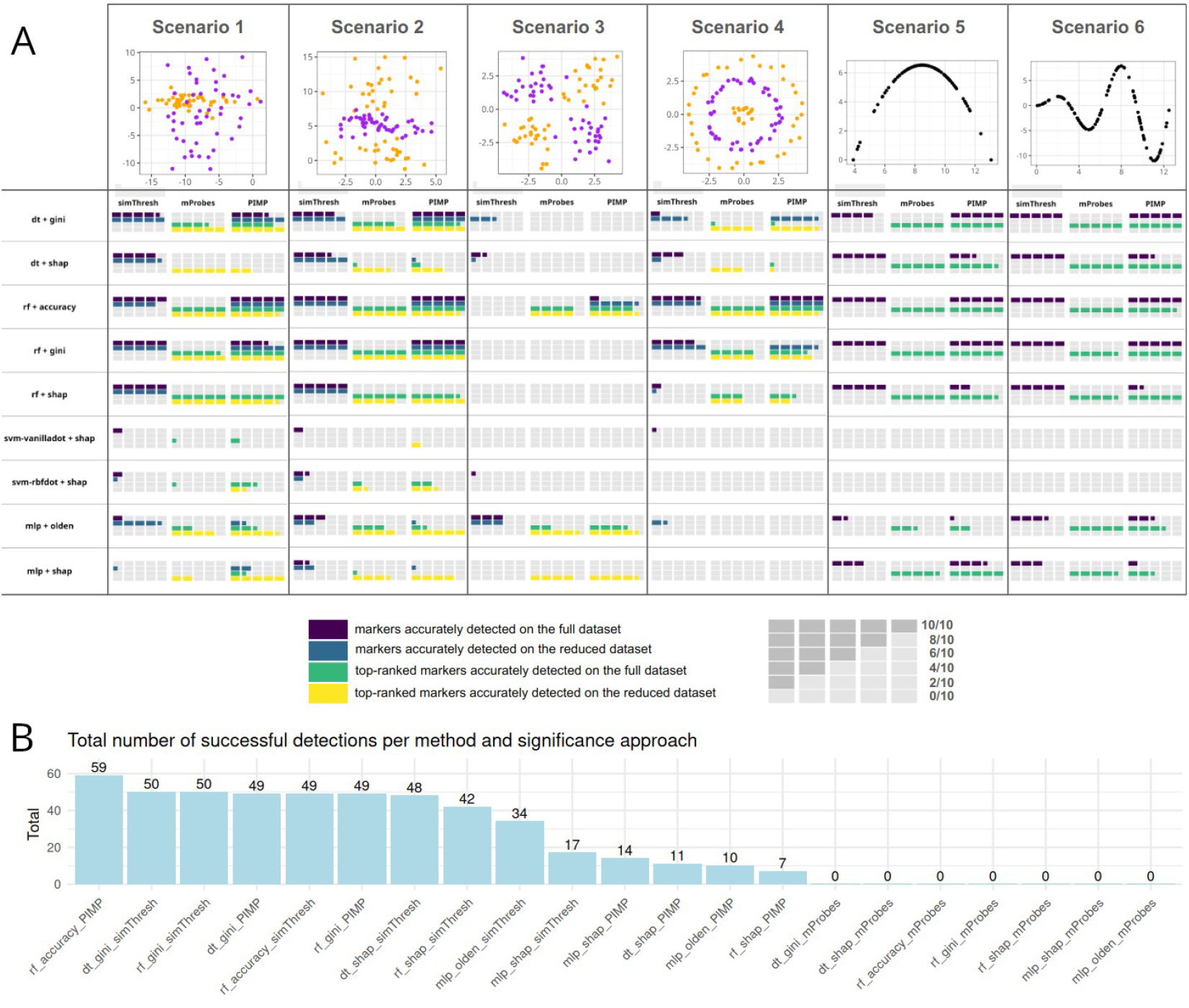
**A**. Variable-selection performance across six simulated scenarios. Columns correspond to Scenarios 1–6. Rows list the model + feature-importance pair. For each scenario, three blocks report the significance procedure (simThresh, mProbes, PIMP). Within each block, colored tiles count successes over 10 replicates for four criteria: exact recovery on the full dataset 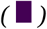, exact recovery on the reduced dataset 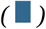, top-rank recovery on the full dataset 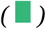, and top-rank recovery on the reduced dataset 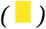. The number of colored tiles equals the number of successful replicates (0–10). This layout enables side-by-side comparison of workflows across scenarios and significance strategies **B**. Total number of detected scenario replicates for each model/feature importance/significance combination.

Across these approaches, the combination RF + accuracy + PIMP achieved the best overall performance, detecting 59 out of 60 relevant features across scenarios and replicates. It was followed by RF or DT with Gini importance and simThresh, which both achieved 50 detections out of 60. Considering only methods with detection rates above 50% (i.e., more than 30 detections), simThresh appeared six times and PIMP three times, highlighting the strong overall contribution of simThresh-based strategies to robust feature identification. However, the gain in detection power brought by PIMP comes at a significant computational cost: indeed, the RF + accuracy + PIMP combination requires roughly 50 times more runtime than tree-based models with Gini importance and simThresh (Fig. 3). Therefore, the choice between these two approaches should be guided by practical considerations such as dataset size and available computational resources, with RF + accuracy + PIMP being preferable when maximizing detection performance is the priority, and DT/RF + Gini + simThresh offering a more scalable alternative for large omics datasets.

## Conclusion

In this work, we benchmarked a wide range of XAI strategies for feature selection across six challenging synthetic scenarios designed to mimic real-world data complexities. By combining different machine learning models, feature importance algorithms, and statistical significance methods, we were able to quantify their respective strengths, limitations, and trade-offs in terms of detection power and computational efficiency.

Our results show that no single combination is universally optimal. The RF + accuracy + PIMP approach achieved the highest overall detection performance, recovering 59 out of 60 cases, but at a substantial computational cost, which may limit its applicability to very large datasets. In contrast, DT or RF with Gini importance and simThresh offered lower detection power but remained highly efficient and scalable, making them strong candidates for exploratory analyses or large-scale studies.

We also found that native feature importance measures (Gini, accuracy, Olden) consistently outperformed SHAP in most scenarios, despite SHAP’s model-agnostic appeal, and that simThresh emerged as the most effective significance approach in the majority of tested conditions.

There is still room for improving XAI-based variable selection, particularly for scenarios involving complex relationships between features, which remain challenging for most pipelines. The *XAItest* framework provides a practical foundation for this progress: it enables easy regeneration of scenario variants and facilitates rapid testing of new model + importance + significance combinations, making it well suited for future methodological development.

Together, these findings provide practical guidance for researchers seeking to integrate XAI into biomarker discovery workflows. They highlight the importance of carefully balancing detection accuracy and computational resources, and they demonstrate how combining interpretable models with appropriate feature importance and significance methods can significantly improve variable selection in complex, high-dimensional omics datasets.

## Code availability

### Bioconductor package XAItest

https://www.bioconductor.org/packages/devel/bioc/html/XAItest.html

### Scenario datasets

https://www.dropbox.com/scl/fo/sg9e1zvu385ovhwf6dp4w/AAWSDwDdiAgvJj2rlq9cbQk?rlkey=2g0ntq8errn0x26wnpm11id6j&st=ukvhtm1h&dl=0

### Paper workflow computations

https://www.dropbox.com/scl/fi/4kjklhh0tfwaa4d1da2yy/0006_R_run10X_simData-F.ipynb?rlkey=a0qij6p7xxkgcs6su5mgk0amb&st=b68xu398&dl=0

### Results analysis

https://www.dropbox.com/scl/fi/902h33wbnuo669dqzpkhc/0009_R_results_tableX10.ipynb?rlkey=61y4ryvfbg9nmf7lzc63rkqn3&st=dk2au0sa&dl=0

https://www.dropbox.com/scl/fi/px9qbtdsdfd61xscde7hk/0010_R_analysis_results_tableX10.ipnb?rlkey=9f93ijlw2x0nnpjzdvbf0oxcf&st=cdgfhdqv&dl=0

## Supplementary Data

### Supp table 1

https://www.dropbox.com/scl/fi/lmyq6n4tnpesslftr760u/supp01.txt?rlkey=n96eqi4l9cza4alv6y1t6oyss&st=5z9j6850&dl=0

### Supp table 2

https://www.dropbox.com/scl/fi/fnlzq7pfggla61vfswsna/supp02.txt?rlkey=mqxsm9heffzmu1xvyhqzocn6v&st=3khtacj4&dl=0

